# Modulating populational variance of methyl-guanine methyl transferase expression through miR-181d degradation: a novel mechanism of temozolomide resistance

**DOI:** 10.1101/2025.04.22.650094

**Authors:** Gatikrushna Singh, Shilpi Singh, Iteeshree Mohapatra, Stefan Kim, Mayur Sharma, Johnny Akers, Thien Nguyen, Eric Wong, Margot Martinez Moreno, Efrosini Kokkoli, Shobha Vasudevan, Sean E. Lawler, Wafik S. El-Deiry, Ziya Gokaslan, Clark C. Chen

**Author notes:** Corresponding Author: Gatikrushna Singh, PhD Department of Neurosurgery University of Minnesota, 515 Delaware St. SE Moos 2-153 Minneapolis, MN-55455. The authors contributed equally as co-senior authors (Gatikrushna Singh, Clark C. Chen). **Author Contributions:** G.S. and C.C.C. designed research; S.S., I.M. and G.S. performed research; S.K., M.S., J.A., T. N., E. W., M.M.M. contributed new reagents/analytic tools; S.S, I.M., E.K., S.V., S.E.L., W.S.D., Z.G, G.S. and C.C.C. analyze data; G.S. and C.C.C. supervised the research; G.S. and C.C.C. wrote the paper. **Competing Interest Statement:** The authors declare no competing interest. **Classification:** Biological Sciences.

## Abstract

Intratumoral heterogeneity plays a pivotal role in cancer evolution, providing the substrate for adaptation to selective pressures, including treatment with chemotherapy. Here, we show that micro-RNA regulation of variance in the expression of the DNA repair protein methyl-guanine methyl transferase (MGMT) contributes to this heterogeneity and acquired therapeutic resistance. In cell lines derived from glioblastomas, the most common form of primary brain tumor, treatment with standard-of-care temozolomide chemotherapy triggers a feed-forward loop between polyribonucleotide nucleotidyltransferase 1 (PNPT1) and miR-181d, an MGMT regulating miRNA, expediting miR-181d degradation. This degradation requires the activation of Ataxia Telangiectasia and Rad3-related (ATR) kinase. The degradation of miR-181d in glioblastoma cells increased both the mean and the variance of MGMT expression in the cell population. Subclone reconstituted cell populations with similar populational mean MGMT levels but with differences in the variance of MGMT expression exhibited differential temozolomide sensitivity, with the higher MGMT variance population showing increased resistance. This resistance is suppressed by exogenously transfected miR-181d. These findings suggest a key role for miRNA in regulating intra-tumoral heterogeneity through modulation of key DNA repair enzymes and provide a compelling rationale for miRNA delivery as a platform for glioblastoma therapy.

**Significance Statement:** This study demonstrates a mechanistic link between a feed-forward loop mediating microRNA degradation and cell-to-cell variance in gene expression, and the contribution of this mechanism to intratumoral heterogeneity and therapeutic resistance. We show that when glioblastoma, the most common form of adult primary brain tumor, is treated with standard-of-care chemotherapy, temozolomide, a feed-forward loop between miR-181d and PNPT1 is initiated, causing rapid degradation of miR-181d. This degradation increases the cell-to-cell variability in methyl-guanine methyl transferase (MGMT) expression, expanding intra-tumoral heterogeneity and contributing to acquired temozolomide resistance. This process can be suppressed by therapeutic delivery of microRNA, providing compelling considerations for clinical translation.

## Introduction

The concepts of mean and variance serve complementary roles in biological research. While the former captures the central tendencies of a trait in populations, the latter defines the degree of spread or dispersion of the data points relative to that central tendency (1). Studies that aim to compare population differences in particular phenotypes (e.g., the expression of a protein before and after a perturbation) often emphasize the differences in means, using variance as a mathematical tool to interpret such differences (2). However, the concept of variance holds independent biological meanings beyond this role. Variance is a key indicator of the complexity of the population and measures differences between the units of the population (e.g. cell-to-cell variability) (3) and is a key determinant of intra-tumoral heterogeneity. Increasing gene expression variance within a population enhances phenotypic diversity to improve the overall fitness of a cell population in response to selective pressures, such as treatment of tumor cells with chemotherapy.

MicroRNAs (miRNAs) are a class of small (19-24 bp), non-coding RNAs that play two key roles in modulating gene expression at the post-transcriptional level (4). miRNAs bind to complementary sequences on target mRNAs, which can accelerate their degradation and/or inhibition of translation (5, 6). These effects ultimately reduce the mean, steady-state expression of the target mRNA in a cell population. Because both RNA degradation and translational inhibition affect stochastic bursts in transcription (7, 8), miRNA additionally decreases the variance in gene expression (9–11). While such suppression enhances the robustness of a cell in maintaining constancy in gene expression, it also reduces the fitness of the population by suppressing transcriptional diversity (12). The effect on population fitness can be magnified when miRNA-regulated genes play central roles in survival in response to selective pressure, such as DNA repair genes in response to DNA-damaging chemotherapeutic agents (13–15).

The DNA repair protein, O^6^ methyl-guanine methyl transferase (MGMT), encodes an evolutionarily conserved DNA repair protein that confers resistance to DNA alkylating agents. Temozolomide (TMZ) is a DNA alkylating agent that serves as the standard-of-care chemotherapy for glioblastoma, the most common form of primary brain cancer in adults. TMZ exerts its anti-tumoral effect predominantly by alkylating the O^6^ position of guanine (16, 17). The presence of O^6^-modified guanine in DNA induces DNA replication arrest and accumulation of single-stranded DNA (ssDNA) breaks (18), triggering DNA damage response (DDR) through activation of the Ataxia Telangiectasia and Rad3-related (ATR) kinase (19). MGMT restores O^6^ alkylated guanine to undamaged guanine (20, 21) by transferring the alkyl group to itself. Each MGMT molecule serves as an acceptor for a single alkyl transfer (16, 22). Thus, high levels of MGMT are associated with glioblastoma resistance to TMZ (23, 24). Of note, MGMT is post-transcriptional regulated by a network of microRNAs (miRNAs); principle amongst these regulators is miR-181d (25, 26). In clinical glioblastoma specimens, high levels of miR-181d were associated with low MGMT expression and improved survival after TMZ treatment (26, 27).

A steady-state level of miRNA is determined by an equilibrium between biogenesis and degradation (28). Relative to the available literature on miRNA biogenesis (29–31), studies of miRNA degradation remain sparse (32–34). The human polyribonucleotide nucleotidyltransferase 1 (PNPT1) encodes a phosphorylase essential for the degradation of select miRNAs (35). The encoded protein belongs to an evolutionarily conserved family of 3’-5’ exo-ribonucleases that use phosphate to catalyze RNA degradation (36, 37). All PNPase family members harbor two RNase PH domains (RPH), an S1 RNA binding domain, and a K-homology domain (37). The crystal structure of bacterial PNPases revealed that the six RPH domains of the PNPase trimer form a doughnut-like structure, with a central channel through which the RNA to be degraded is threaded (38). Beyond miRNA degradation, the human PNPT1 plays a pivotal role in the import of RNA into the mitochondria. Separate-of-function PNPT1 mutations have been reported for these distinct functions (39, 40).

In this study, we demonstrate that TMZ treatment activates the ATR kinase to initiate a feed-forward PNPT1-miR-181d loop that results in the degradation of miR-181d by PNPT1. The decreased miR-181d level triggered two independent mechanisms contributing to acquired TMZ resistance: 1) increasing the population mean expression of MGMT and 2) broadening the cell-to-cell variability in MGMT expression. Over-expression of miR-181d suppressed both resistance mechanisms and synergized with TMZ in tumoricidal activity against glioblastoma. These findings provide a compelling rationale for miRNA delivery-based therapy to suppress chemotherapy-induced genetic heterogeneity that underlies therapeutic resistance.

## Results

### Decreased miR-181d expression following temozolomide treatment

To characterize the changes in miRNA expression in response to TMZ therapy, two patient-derived glioblastoma cell lines BT-83 (41) and CMK3 (42) were treated with 500 µM TMZ or vehicle for 6 h. These cell lines were selected because their gene expression profile suggests they capture the two key glioblastoma cell states: BT-83 exhibits an expression pattern suggestive of the mesenchymal-like state, while CMK3 exhibits an expression pattern suggestive of the neural progenitor-like state (43). RNA was extracted and profiled by RNA sequencing (44). In both cell lines, the TMZ treatment did not significantly alter the expression of >95% of the miRNAs (**Fig. 1A, Supp Fig. 1A**). miRNAs exhibiting a >2-fold change in expression following TMZ treatment in both lines were individually assessed to determine whether their expression differed in matched pre- and post-TMZ treated clinical glioblastoma specimens (**Fig 1B, Supp Fig. 1B**. Among the miRNAs evaluated, miR-181d was the only one consistently showing lowered levels in the post-TMZ specimens (**Fig. 1B**). This finding recapitulated our prior profiling experiment, where miR-181d was also observed to be downregulated in post-TMZ treated clinical glioblastoma samples (45).

**Figure 1.**
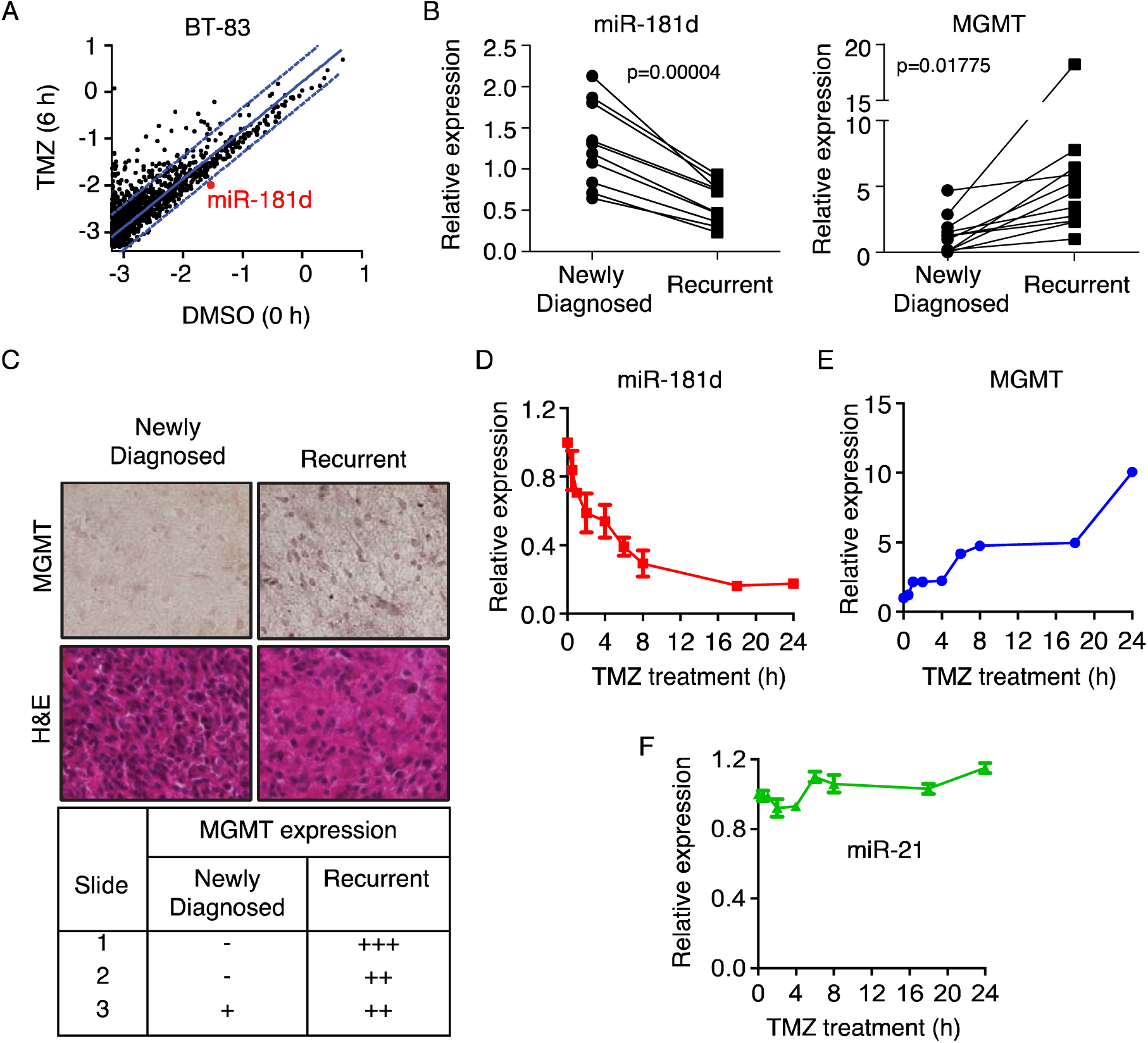
TMZ induces degradation of miR-181d and upregulation of MGMT. **(A)** Scatter plot representing the differential expression analysis of miRNAs from RNA-seq data of patient-derived BT-83 glioblastoma cells treated with 500 µM TMZ for 6 h. miRNA expression levels in TMZ-treated cells were compared to vehicle-treated cells (DMSO). miRNAs falling outside the diagonal lines indicate significant expression changes following TMZ treatment. miR-181d, which shows a notable reduction, is highlighted in red. **(B)** Quantification of miR-181d and MGMT expression in 10 matched pairs of newly diagnosed (pre-TMZ treatment) and recurrent (post-TMZ treatment) formalin-fixed paraffin-embedded (FFPE) glioblastoma surgical specimen. miR-181d and MGMT expression were measured using RT-qPCR. **(C)** Immunohistochemistry (IHC) analysis of MGMT in paraffin-embedded human glioblastoma specimens from (B). MGMT staining intensity was scored and summarized in the table below. Hematoxylin and Eosin (H&E) staining represents the even distribution of cells in both newly diagnosed and recurrent glioblastoma specimen sections (4 μm). MGMT expression: No (-), Low (+), Intermediate (++), High (+++). **(D, E, F)** Patient-derived CMK3 glioblastoma cells were treated with 500 µM TMZ up to 24 h. RNA was isolated and analyzed by RT-qPCR. miRNA and MGMT expression were normalized to vehicle-treated (DMSO) controls. miR-181d (**D**), MGMT (**E**) and miR-21 (**F**).

Since miR-181d down-regulates MGMT expression (25, 27), we next characterized the MGMT mRNA expression in the matched pre- and post-TMZ treated clinical glioblastoma specimen by qPCR. The lowered levels of miR-181d in the post-TMZ treated specimens were associated with increased levels of MGMT expression (**Fig. 1B**). MGMT immunohistochemical staining was also performed using formalin-fixed paraffin-embedded (FFPE) sections derived from three pairs of matched newly diagnosed and recurrent clinical glioblastoma specimens. Increased MGMT expression was consistently observed in the recurrent tumor sections relative to the newly diagnosed tumor sections in all three matched pairs (**Fig. 1C**).

To characterize the effect of TMZ treatment on miR-181d expression, the CMK3 glioblastoma line was treated with TMZ and analyzed by RT-qPCR. This analysis revealed a time-dependent reduction in miR-181d levels (**Fig. 1D**), with >50% decrease in miR-181d was observed within two hours of TMZ treatment. This decrease in miR-181d level was associated with a concomitant increase in MGMT mRNA expression (**Fig. 1E**). As a control, the expression of another miRNA, miR-21, remained unaffected by TMZ treatment (**Fig. 1F**).

### Polyribonucleotide nucleotidyltransferase 1 (PNPT1) is required for miR-181d degradation

A decrease in the steady-state level of miR-181d following TMZ treatment could result from either suppressed biogenesis or increased degradation. To distinguish between these possibilities, the effect of TMZ on both precursor (pre) and mature (mat) miR-181d was examined. The level of pre-miR-181d remained unchanged in response to TMZ treatment in CMK3 cells. In contrast, the level of mat-miR-181d significantly decreased after treatment (**Fig. 2A**). Similar results were observed in the BT-83 line (**Supp. Fig. 2A**). These observations suggest that the reduced level of miR-181d resulted from a degradative process rather than suppressed biogenesis. Supporting this hypothesis, inhibition of transcription by pre-treatment with Actinomycin D1 (46, 47) did not significantly alter the TMZ-induced reduction in miR-181d levels (**Fig. 2B**).

**Figure 2.**
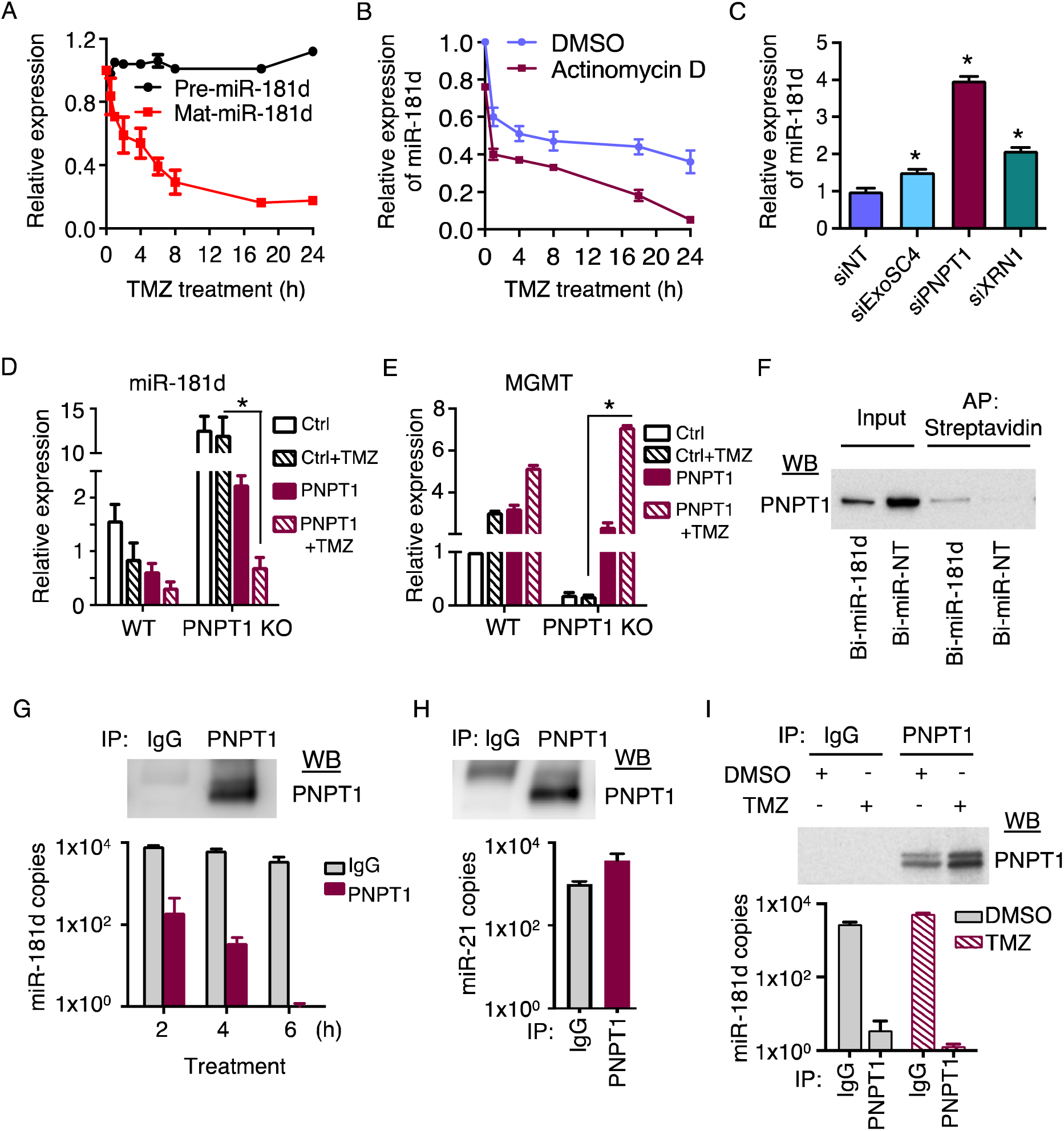
PNPT1 is required for TMZ-induced degradation of miR-181d. **(A)** CMK3 cells were treated with 500 µM TMZ up to 24 h. RNA was isolated, and the precursor (pre) or mature (mat) miR-181d expression was analyzed using RT-qPCR. miRNA expression was normalized to vehicle-treated (DMSO) controls. **(B)** CMK3 cells were pretreated with Actinomycin D1 (100 nM) and exposed to TMZ (500 µM) for 24 h. miR-181d expression was analyzed by RT-qPCR. **(C)** CMK3 cells transfected with siExoSC4, siPNPT1, or siXRN1 for 24 h. miR-181d expression was analyzed by RT-qPCR. Statistical significance: \**p*<0.05. **(D, E)** Overexpression of PNPT1 restored TMZ-induced miR-181d degradation and increased MGMT expression. CMK3 WT or PNPT1 KO cells were transfected with pCMV6-AC-GFP-PNPT1 overexpressing plasmid and exposed to TMZ or DMSO. miR-181d **(D)** or MGMT **(E)** expression were analyzed by RT-qPCR. Statistical significance: \**p*<0.05 between the indicated groups. **(F)** miR-181d interacts with PNPT1. CMK3 cells transfected with biotinylated (Bi)-miR-181d or miR-NT control mimic. Bi-miRNA was affinity purified (AP) with streptavidin-magnetic beads. The associated protein complex was eluted and analyzed by Western blotting. Total cell lysate was used as input. **(G, H)** PNPT1 protein was immunoprecipitated and incubated with miR-181d or miR21 mimic (1X10^5^ copies) for 6 h. RNA was extracted from the reaction mixture and quantified by RT-qPCR. miR-181d **(G)**, miR-21 **(H)**. **(I)** Affinity-purified PNPT1 was isolated from CMK3 cells with and without TMZ treatment and co-incubated with miR-181d mimic.

To better understand the mechanism underlying TMZ-induced miR-181d degradation, a siRNA screen was conducted targeting RNases (ExoSC4, PNPT1, XRN1) previously implicated in miRNA degradation. Among the tested RNase targeting siRNAs, only those directed against PNPT1 (48) caused an increase in the steady-state miR-181d levels (a four-fold increase, **Fig. 2C**). This effect was confirmed using CRISPR/Cas9 disruption of PNPT1, which caused a> 100-fold increase in miR-181d (**Supp. Fig. 2B, C**). PNPT1 knockout (KO) also prevented the TMZ-induced reduction in miR-181d levels (Supp. **Fig. 2D**).

Ectopic overexpression of PNPT1 in both wild type (WT) or PNPT1 KO CMK3 cells reduced the steady-state levels of miR-181d (**Fig. 2D**), with an associated rise in MGMT mRNA expression (**Fig. 2E**). Moreover, such over-expression enhanced TMZ-induced decrease in miR-181d (**Fig. 2D**), with associated increase in MGMT (**Fig. 2E**). To investigate the interaction between miR-181d and PNPT1, CMK3 cells were transfected with biotinylated (Bi)-miR-181d or Bi-miR-NT control mimic. A streptavidin-biotin pull-down assay was performed, followed by PNPT1 detection using western blotting. The results demonstrated that PNPT1 co-precipitated with Bi-miR-181d in the streptavidin pull-down (**Fig. 2F**).

To determine whether PNPT1 is required for miR-181d degradation, an *in vitro* miRNA degradation assay was performed. PNPT1 protein was immunoprecipitated (IP) using an anti-PNPT1 antibody from CMK3 cell lysates (**Fig. 2G**) and incubated with mat-miR-181d or miR-21 mimic. The miRNAs were subsequently isolated from the reaction mixture and analyzed by RT-qPCR. The results demonstrated a time-dependent decrease in miR-181d level (**Fig. 2G**). Such a decrease was not observed with miR-21 mimic (**Fig. 2H**). Moreover, the miR-181d decrease was more pronounced when incubated with affinity-purified PNPT1 derived from CMK3 treated with TMZ than that observed using affinity-purified PNPT derived from CMK3 treated with vehicle (**Fig. 2I**). Together, these findings suggest the involvement of PNPT1 in miR-181d degradation.

### PNPT1 modulates TMZ sensitivity through miR-181d and MGMT regulation

The effect of PNPT1 on TMZ sensitivity was evaluated by the limiting dilution neurosphere forming assays (49, 50). CMK3 cells were transfected with PNPT1 siRNAs and treated with TMZ (500 µM) or vehicle to determine the clonogenic potential. The result demonstrated that PNPT1 silencing (**Supp. Fig. 3A**) caused an approximate two orders of magnitude increase in TMZ sensitivity (**Supp. Fig. 3B** top panel). Similarly, transfection of miR-181d mimic or MGMT targeting siRNA into CMK3 caused an approximate two orders of magnitude increase in TMZ sensitivity (**Supp. Fig. 3B**, middle and bottom panel). siPNPT1-mediated increase in TMZ sensitivity can be blocked by an anti-miR against miR-181d (**Supp. Fig. 3C**, top panel) or MGMT overexpression (**Supp. Fig. 3C**, bottom panel). These results support a genetic pathway where PNPT1 acts on miR-181d to regulate MGMT expression level and modulate TMZ sensitivity.

### Mutations inactivating PNPT1 ribonuclease activity suppressed its effects on miR-181d

Next, we determined the expression of PNPT1 in clinical glioblastoma specimens. PNPT1 expression was elevated by 2 to 25-fold in tumor specimens relative to the adjacent brain tissue at the mRNA (**Fig. 3A**) and protein level (**Fig. 3B**). PNPT1 is a multi-functional protein comprising two RNase PH (RPH) domains, two COOH-terminal RNA binding domains: an S1 RNA binding domain and a K-homology domain (51). Separation-of-function mutations between its two key activities, mitochondrial RNA import, and exoribonuclease activity have been described (39). Specifically, mutations S135G, Q387R, and S484A disrupted PNPT1 mitochondrial function without significantly affecting ribonuclease activity (52–54), while mutations R445E, R446E, D538A, and D544G inactivated exoribonuclease activity (53). To assess the impact of these mutations on miR-181d degradation, expression constructs of PNPT1 WT and mutants were transfected into CMK3 (as control) or two independent clones of PNPT1 KO lines. Expression of WT-PNPT1 in the PNPT1 KO lines suppressed the steady-state miR-181d level by approximately an order of magnitude (**Fig. 3C**). Transfection of PNPT1 constructs bearing the S135G, Q387R, or S484A mutations do not significantly affect the exo-ribonuclease activity, resulted in suppression of steady-state miR-181d level comparable to WT-PNTP1. In contrast, constructs bearing the R445E, R446E, D538A, or D544G mutations (mutations that disrupted the exo-ribonuclease activity) did not significantly affect steady-state miR-181d level (**Fig. 3C**).

**Figure 3.**
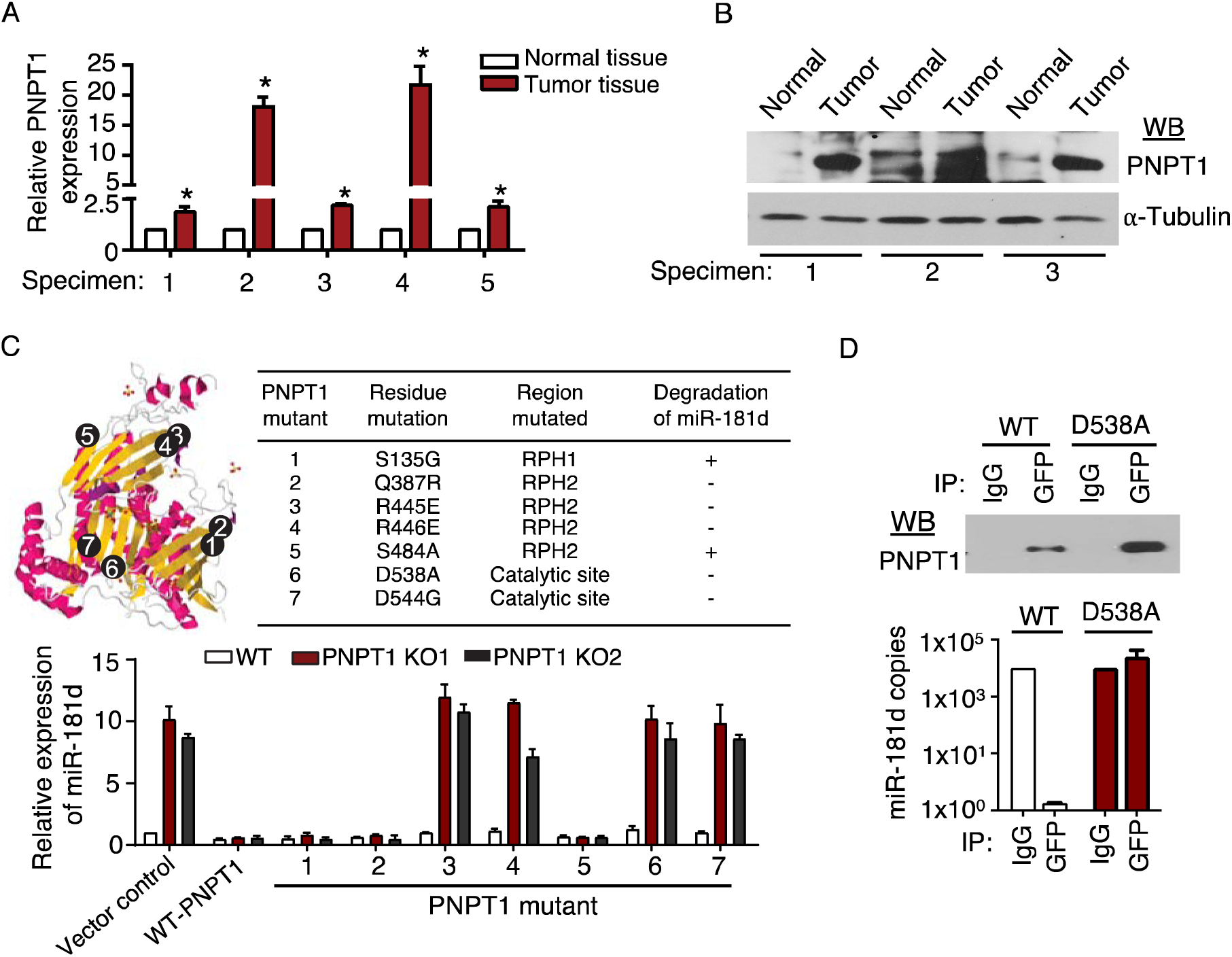
Ribonuclease activity of PNPT1 degradation of miR-181d. PNPT1 expression analysis in clinical glioblastoma specimens. RNAs (n=5) and protein (n=3) extracted from clinical glioblastoma and matched adjacent brain tissue specimens were analyzed for PNPT1. PNPT expression is elevated in tumor tissues relative to adjacent normal brain tissues. **(A)** PNPT1 mRNA expression by RT-qPCR. Statistical significance: *p<0.05. **(B)** PNPT1 protein expression by Western blotting. **(C)** Left panel: PNPT1 protein structure labelled with numbers represents the specific position for the amino acid substitution mutation. Right panel (table): Summary of amino acid substitution within PNPT1 domains. Bottom panel: CMK2 wild-type (WT) and PNPT1 knockout clones (PNPT1 KO1, PNPT1 KO2) were transfected with either WT or mutated PNPT1 constructs. miR-181d expression was quantified by RT-qPCR. **(D)** *In vitro* miR-181d degradation assay by immunoprecipitated WT or mutant PNPT1. GFP-tagged PNPT1 protein was immunoprecipitated using anti-GFP antibody from WT or D538A mutant PNPT1 expressing lines and incubated with miR-181d mimic. RNA was extracted and quantified by RT-qPCR.

To further establish the involvement of PNPT1’s exoribonuclease activity in miR-181d degradation, GFP-tagged WT-PNPT1 or the D538A mutant were affinity-purified from stably expressing CMK3 PNPT1 KO cells and incubated with mature miR-181d mimic. The level of miR-181d remained unchanged when incubated with the PNPT1 D538A, while a significant reduction of miR-181d was observed when incubated with WT-PNPT1 (**Fig. 3D**). These results suggest that PNPT1 exo-ribonuclease activity contributes to the degradation of miR-181d.

### PNPT1 is post-translationally regulated by miR-181d

Sequence analysis of the PNPT1 3′ untranslated region (3′UTR) revealed three potential miRNA Response Elements (MREs) for miR-181d, depicted as block1 (B1:200 bp); block2 (B2: 300 bp); block3 (757 bp) (**Fig. 4A**). This raises the possibility that PNPT1 is regulated by miR-181d in a feed-forward loop, where PNPT1 expression increases following TMZ-induced miR-181d degradation. Supporting this hypothesis, the expression of PNPT1 increased in response to TMZ treatment at both the RNA (**Fig. 4B**) and protein (**Fig. 4C**) levels. Further supporting this hypothesis, increased PNPT1 mRNA expression was observed in matched pre- and post-TMZ-treated clinical glioblastoma specimens (**Fig. 4D**).

**Figure 4.**
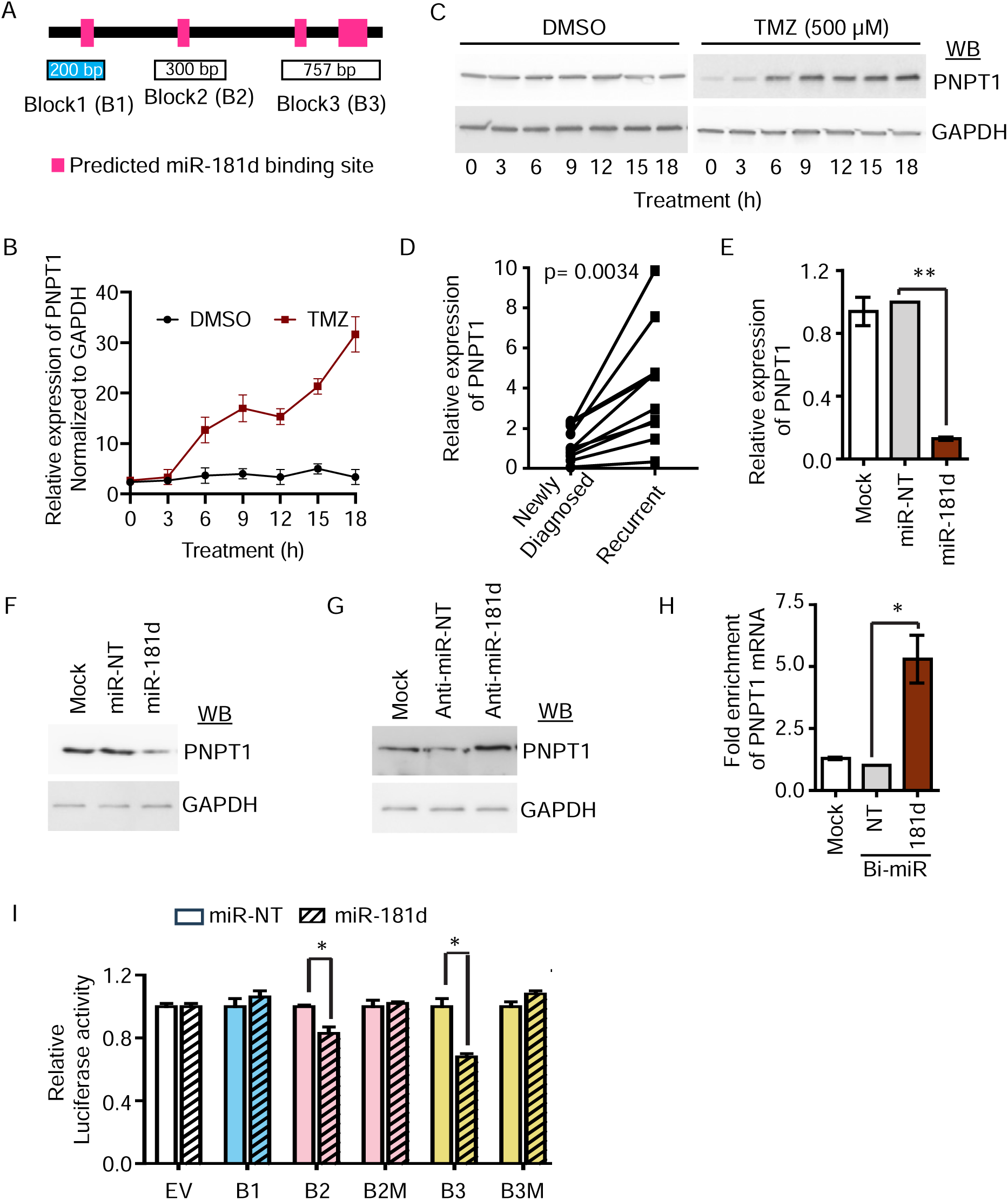
miR-181d regulates PNPT1 expression. **(A)** Schematic representation of miR-181d miRNA Response Elements (MREs) in PNPT1 3’UTR. Pink block: Predicted MREs of miR-181d (Block1: 200 bp, Block2: 300 bp, Block3: 757 bp). **(B)** CMK3 cells were treated with TMZ or DMSO for the indicated time point. Total RNA was extracted, and PNPT1 expression was analyzed using qPCR. GAPDH was used as an internal loading control. **(C)** PNPT1 protein expression by western blotting in CMK3 cells used in (B). **(D)** TMZ treatment increases PNPT1 mRNA in recurrent glioblastoma. PNPT1 mRNA expression was quantified in 10 matched pairs of newly diagnosed and recurrent glioblastoma specimens using RT-qPCR. **(E, F)** miR-181d suppress mRNA and protein expression of PNPT1. CMK3 cells transfected with miR-181d or miR-NT control mimic. PNPT1 mRNA expression was measured by RT-qPCR. **p<0.001 indicates statistically significant differences compared to miR-control **(E).** Protein expression was analyzed by Western blotting **(F)**. **(G)** Anti-miR-181d de-repressed PNPT1 expression. CMK3 cells transfected with anti-miR-181d or anti-miR NT. The cell lysate was analyzed for PNPT1 expression by Western blotting. **(H)** miR-181d mimic binds to PNPT1 mRNA. CMK3 cells transfected with Bi-miR181d or NT control mimic. The lysate was affinity purified with streptavidin magnetic beads. Isolated RNA was analyzed for PNPT1 by RT-qPCR. Statistical significance *p<0.05. **(I)** Luciferase reporter assay of miR-181d MREs and its mutants in PNPT1 3’UTR with miR-181d or NT control mimic. Statistical significance *p<0.05.

To further substantiate miR-181d regulation of PNPT1, CMK3 cells were transfected with human miR-181d or miR-NT control mimic. Forty-eight hours post-transfection, RT-qPCR detected ∼80% reduction in PNPT1 mRNA (**Fig. 4E**) and protein expression (**Fig. 4F**). In addition, transfection of anti-miR-181d into CMK3 cells that expressed high levels of miR-181d, increased PNPT1 protein expression. This effect was not observed after the transfection of a control anti-miR-NT (**Fig. 4G**). Next, we determined whether PNPT1 mRNA binds to miR-181d. CMK3 cells were transfected with Bi-miR-181d or Bi-miR-NT control mimic. A streptavidin-biotin pull-down assay was performed, followed by the detection of PNPT1 mRNA by RT-qPCR. The analyses detected approximate 5-fold enrichment of PNPT1 mRNA with the Bi-miR-181d pull-down relative to Bi-miR-NT control pull-down (**Fig. 4H**).

To further confirm that PNPT1 is a target of miR-181d, each of the three miR-181d MREs was cloned into a pSiCheck-2 reporter construct. Luciferase activity of a pSiCheck-2 reporter construct bearing the first miR-181d MRE (B1) was unaffected by the introduction of miR-181d. However, the construct harboring the second (B2) and third (B3) miR-181d MREs exhibited ∼20% and ∼30% reductions in luciferase activity, respectively, upon introduction of miR-181d (**Fig. 4I**). Mutations in these MRE sites (B2M and B3M) that disrupted complementarity to miR-181d abolished these effects. These findings suggest that miR-181d modulates PNPT1 expression through specific MREs in its 3′UTR.

### ATR is required for TMZ-induced miR-181d degradation

TMZ treatment initiates DNA damage response (DDR) by activating the Ataxia Telangiectasia and Rad3-related (ATR) kinase (55). To investigate whether ATR activation is necessary for TMZ-induced miR-181d degradation, CMK3 cells were pretreated with the ATR inhibitor VE-821(56), followed by TMZ exposure, and miR-181d expression levels were assessed. While TMZ treatment induced a time-dependent decrease in miR-181d, this reduction was suppressed by VE-821 (**Fig. 5A**).

**Figure 5.**
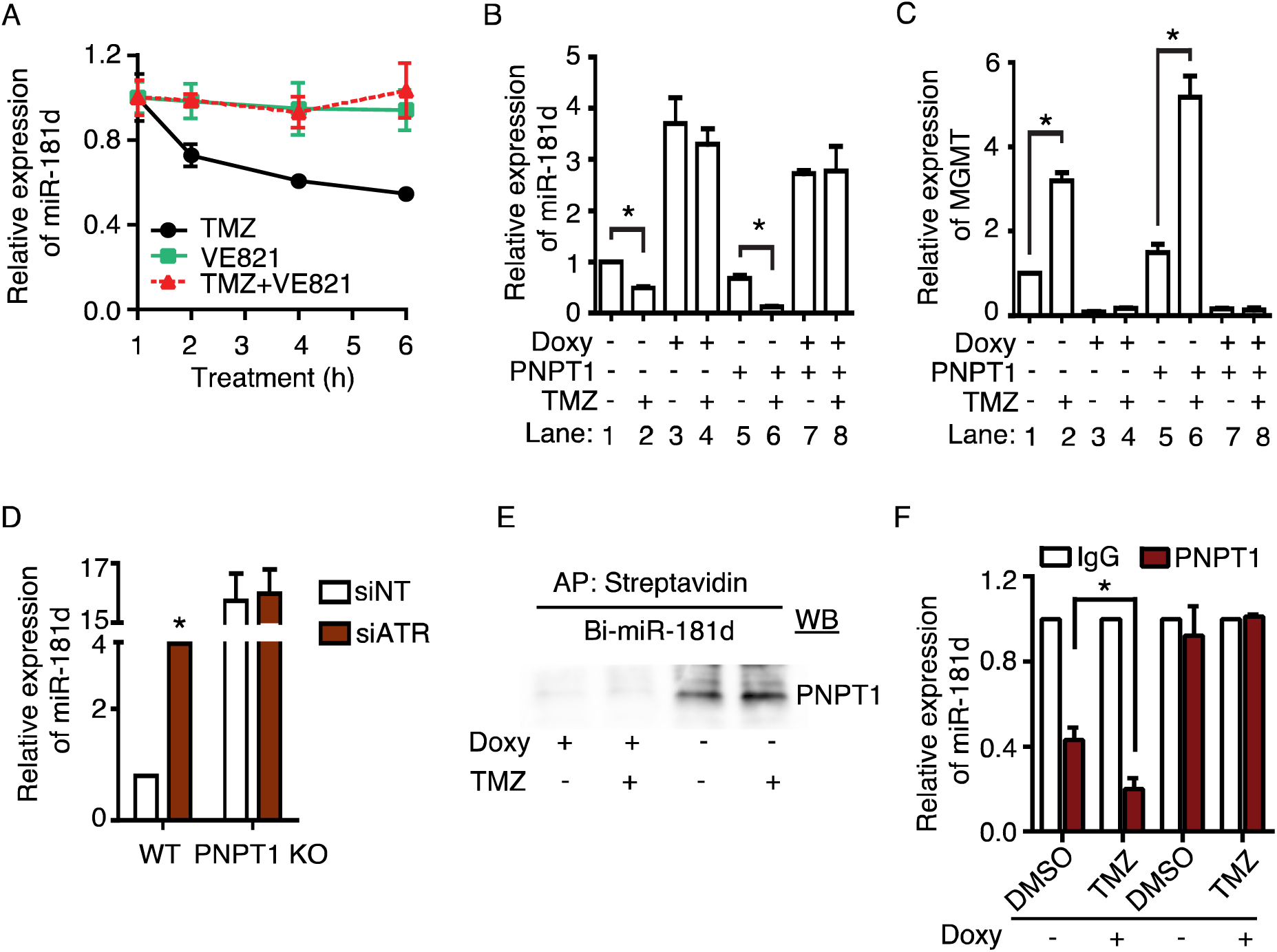
PNPT1 required ATR for TMZ-induced miR-181d degradation. **(A)** The ATR inhibitor, VE-821, suppresses TMZ-induced miR-181d degradation. CMK3 cells were pretreated with ATR inhibitor (VE-821, 10 μM, 12 h) before TMZ (500 µM) treatment. miR-181d level was measured at indicated time points using RT-PCR. **(B, C)** ATR activation is the upstream event of PNPT1 action. Doxycycline (Doxy)-inducible shATR-expressing with or without PNPT1 overexpression in CMK3 cells were treated with TMZ (500 µM) or DMSO. miR-181d **(B)** and MGMT **(C)** expression were analyzed by RT-qPCR. **(D)** Downregulation of ATR elevated miR-181d expression. siATR or siNT was transfected to CMK3 WT or PNPT1 KO cells. miR-181d expression was analyzed by RT-qPCR. **(E)** PNPT1 required ATR for the interaction with miR-181d. Doxycycline (Doxy)-induced shATR were transfected with Bi-miR-181d mimic. Cell lysates were affinity purified with streptavidin and analyzed by Western blotting. **(F)** ATR silencing abolished PNPT1 degradation of miR-181d. shATR induced CMK3 cells treated with TMZ or DMSO. PNPT1 was immunoprecipitated from the cell lysate and incubated with miR-181d mimic for degradation assay. miR-181d was quantified by RT-qPCR.

Next, we tested the interaction between ATR and PNPT1 in their regulation of miR-181d. To this end, we generated a CMK3 glioblastoma line harboring the doxycycline-inducible ATR-silencing (shATR) construct that over-expresses PNPT1. ATR silencing suppressed TMZ-induced miR-181d degradation (**Fig. 5B**, lanes 4 versus lane 3), while PNPT1 over-expression enhanced miR-181d degradation phenotype (**Fig. 5B**, lanes 6 versus lane 2). Notably, ATR silencing in the cell line that over-expressed PNPT1 showed no evidence of TMZ-induced miR-181d degradation (**Fig. 5B**, lane 8 versus 7), suggesting that ATR is required for PNPT1 mediated miR-181d degradation, with associated increased MGMT expression (**Fig. 5C**). To further evaluate the genetic interaction between PNPT1 and ATR, CMK3 or CMK3 PNPT1 KO lines were transfected with siATR or siNT control. In CMK3 cells, siATR transfection resulted in an order of magnitude increase in miR-181d than in siNT transfected cells, while such difference was not observed in the PNPT1 KO line (**Fig. 5D**). These results suggest that the effect of ATR on miR-181d requires PNPT1.

To determine whether ATR is required for binding of PNPT1 to miR-181d, the CMK3 line harboring the doxycycline-inducible shATR was transfected with Bi-miR-181d or Bi-NT and treated with TMZ or vehicle. Streptavidin pull-down was performed using lysate derived from doxycycline-treated or control-treated cells, followed by Western blotting with an anti-PNPT1 antibody. Without doxycycline induction of ATR silencing, PNPT1 was detected in the Bi-miR-181d pull-down, with or without TMZ treatment (**Fig. 5E**). However, upon doxycycline treatment induced ATR silencing, PNPT1 was not detected in the Bi-miR-181d pull-down (**Fig. 5E**), suggesting that ATR is required for PNPT1-miR-181d interaction.

To determine whether ATR is essential for PNPT1-mediated degradation of miR-181d, PNPT1 protein was immunoprecipitated (IP) from doxycycline or vehicle-treated CMK3 harboring doxycycline-inducible shATR subjected to TMZ or DMSO treatment. Lowered level of miR-181d was observed after incubation with affinity-purified PNPT1 relative to reactions incubated with IgG pull-down (**Fig. 5F**, lane 2 versus 1). This effect was enhanced when miR-181d was incubated with PNPT1 affinity purified from TMZ-treated cells (**Fig. 5F**, lane 4 versus 2). No such decrease in miR-181d was observed when miR-181d was incubated with affinity-purified PNPT1 from doxycycline-treated cells, suggesting that ATR is required for PNPT1-mediated degradation of miR-181d (**Fig. 5F**, lanes 4 and 8).

### miR-181d degradation enhanced cell-to-cell variability and temozolomide resistance

miRNAs suppress transcriptional variance in gene expression (57). To determine whether this suppression influences cell-to-cell variability, we isolated individual clones of BT-83 glioblastoma cells following TMZ or vehicle treatment. Single-cell qPCR was performed on approximately 80 cells per treatment group to assess miR-181d and MGMT expression levels, with the data presented as a distribution. Consistent with our hypothesis that TMZ induces degradation of miR-181d, the population mean of miR-181d expression is reduced after TMZ treatment (**Fig. 6A**, top panel). This reduction is associated with an increased population mean of MGMT expression (**Fig. 6A**, bottom panel). Notably, while the variance in miR-181d expression remained unchanged in response to TMZ, the variance of MGMT increased by a factor of five after TMZ treatment (p =0.001). Transfection of a control miRNA did not alter the distributional mean or variance of miR-181d expression (**Fig. 6B**, top panel). On the other hand, transfection of miR-181d mimic increased the distributional mean of miR-181d expression without significantly changing its variance (**Fig. 6B**, top panel). In terms of MGMT expression, miR-181d transfection reduced the distributional mean without affecting the variance of MGMT expression (**Fig. 6B**, bottom panel). Together, these results suggest that miR-181d modulates the populational variance of MGMT expression.

**Figure 6.**
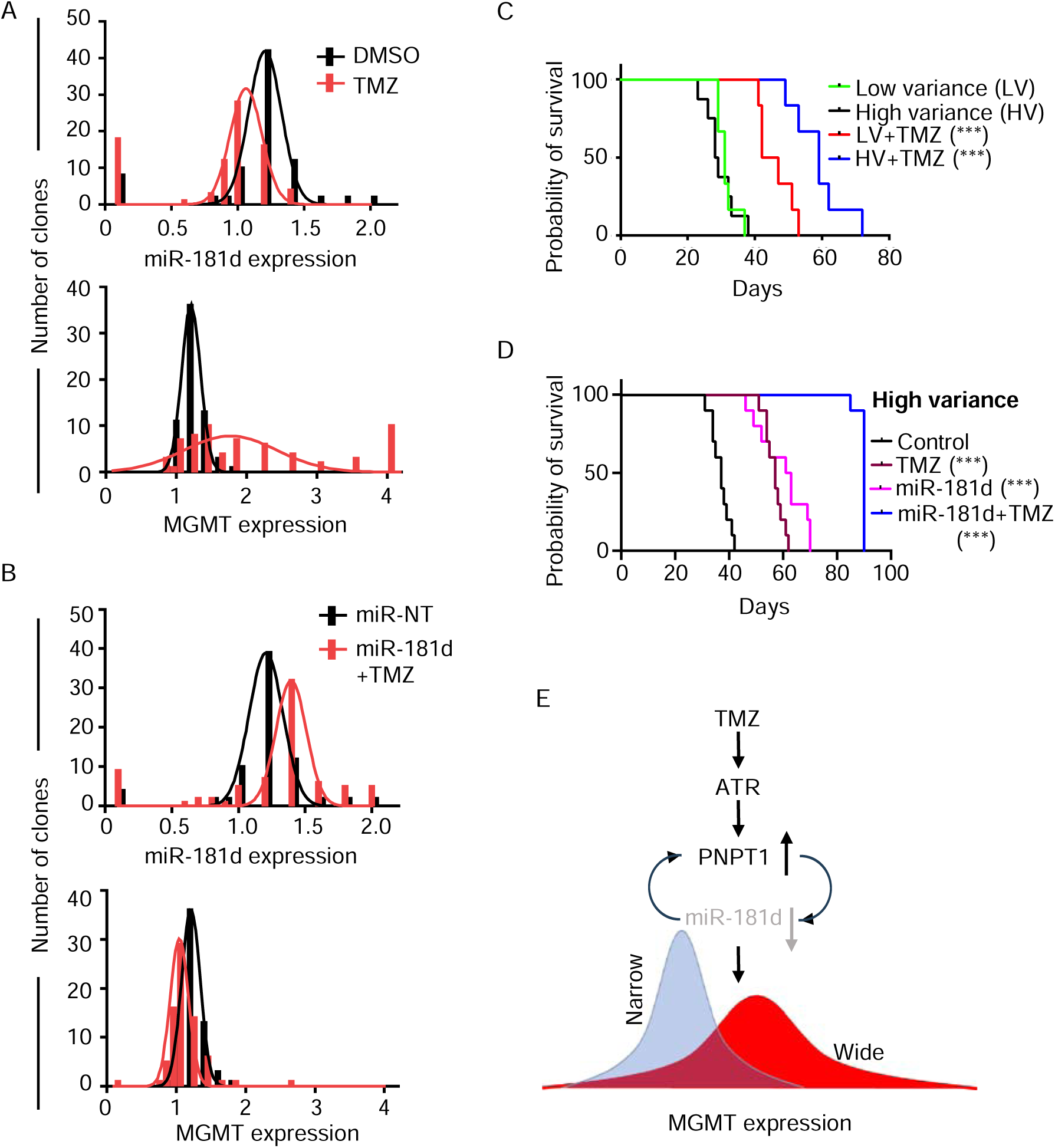
miR-181d mediated cell-to-cell variation of MGMT increase TMZ resistance. **(A)** TMZ reduces miR-181d expression and variation in MGMT expression. BT-83 cells were treated with 500 µM TMZ or DMSO for individual clone generation, RNA was isolated from the individual clones, and RT-qPCR for miR-181d and MGMT expression. **(B)** miR-181d reduced the MGMT expression variation by TMZ. The cells were transfected with miR-181d before TMZ treatment. A single cell of the individual clones was analyzed for miR181d or MGMT expression by RT-qPCR. **(C)** Kaplan–Meier survival curves of nude mice bearing intracranial BT-83 cells selected for low variance (LV) and high variance (HV) in MGMT expression. The mice underwent the intraperitoneal administration of TMZ at 50 mg/kg/day for 5-days followed by a 23-day treatment interruption. TMZ treatment was initiated 7-days post-tumor implantation. Each group consisted of 10 mice. **(D)** The high-variance cells were transfected with miR-181d and implanted into nude mice. The mice were treated with TMZ as described (C). The study was continued for 90 days. **(E)** TMZ treatment of glioblastoma activates ATR and PNPT1, triggering the degradation of miR-181d. Loss of miR-181d, stabilizes PNPT1 expression, creating a feed-forward loop that enhances TMZ-induced miR-181d degradation. This led to an increase in the mean expression of MGMT and greater cell-to-cell variability in MGMT expression.

Next, we tested whether variability in MGMT expression translates to altered TMZ resistance. To this end, we isolated BT-83 subclones that varied in MGMT expression and pooled these subclones to generate pooled populations with comparable MGMT expression but with either low-variance or high-variance in MGMT expression. For the low-variance group, we combined seven clones exhibiting similar MGMT expression levels (**Supp. Fig. 4A**). The high-variance group was created by pooling three clones with MGMT expression levels comparable to the low-variance group, along with two clones exhibiting higher and two clones with lower MGMT expression (**Supp. Fig. 4A)**. Quantitative RT-PCR analysis confirmed that the mean MGMT mRNA expression levels were comparable between the low- and high-variance populations (**Supp. Fig. 4B**). However, Western blotting shows that MGMT protein expression is slightly higher in the low-variance group (**Supp. Fig. 4C**). These two populations were orthotopically implanted into nude mice. The mice were treated with TMZ to assess the impact of MGMT expression variance on therapeutic response.

Without TMZ treatment, the mice bearing low-variance and high-variance MGMT expression pools exhibited similar median survival of 38 and 37 days, respectively (**Fig. 6C**). After temozolomide treatment, the median survival for mice implanted with the low-variance MGMT expression pool extended to 53 days. In contrast, those implanted with the high-variance pool demonstrated a significantly longer median survival of 72 days (p <0.0001), indicating increased resistance to TMZ in the high-variance group (hence prolonged survival) relative to the low-variance MGMT pool. This finding is notable given that the populational MGMT expression level in the low-variance group is slightly higher than that of the high-variance group (**Supp. Fig. 4C**) supporting a role for MGMT expression variance in TMZ resistance.

We next determined whether miR-181d transfection could enhance TMZ sensitivity. The high variance of MGMT-expressing population was transfected with miR-181d before orthotopic implantation. Mice implanted with the high variance pool transfected with miR-181d had a median survival of 70 days, significantly longer than those mice transfected with control miRNA (42 days, p <0.0001, **Fig. 6D**). TMZ treatment of those mice transfected with control miRNA increased median survival to 61 days (p<0.0001). Notably, TMZ treatment in the miR-181d-transfected group further prolonged survival, with all but one mouse surviving the planned study endpoint of 90 days (p<0.0001 compared to TMZ-treated mice implanted with the control miRNA-transfected tumor). These results demonstrate that miR-181d enhanced the tumoricidal effects of TMZ against a glioblastoma population exhibiting cell-to-cell variation in MGMT expression.

## Discussion

Transcriptional variability contributes to phenotypic diversity that underlies populational fitness to selective pressures, such as exposure of cancer cells to chemotherapy (58). Micro-RNAs suppress such transcriptional variability (11, 59, 60). Results from this study demonstrate that DNA-damaging chemotherapy treatment triggers feed-forward loops to mediate micro-RNA degradation, enhancing cell-to-cell variability in the expression of critical DNA repair proteins. Specifically, treatment of glioblastoma cells with the chemotherapy, temozolomide, triggers an ATR- (61) and PNPT1-dependent degradation of miR-181d. Since PNPT1 is itself a target of miR-181d, miR-181d degradation de-repress PNPT1, leading to a feed-forward loop that amplifies TMZ-induced miR-181d degradation, resulting in both 1) an increased mean expression of MGMT as well as the 2) cell-to-cell variability of MGMT expression (**Fig. 6E**). The importance of the former to TMZ resistance is well-established (62). Our study demonstrates that the latter also contributes to acquired TMZ resistance. This result is intuitive since the higher the MGMT-expressing clones in the high-variance pool are more likely to survive TMZ treatment. Moreover, our results indicate that both mechanisms of temozolomide resistance (increased mean and variance of MGMT expression) can be suppressed by miR-181d over-expression, suggesting potential as a therapeutic strategy.

The varied levels of MGMT expression in glioblastoma cells bear implications on intra-tumoral heterogeneity (15). MGMT restores temozolomide-induced O^6^ methylated guanine that accumulates due to the alkylating activities associated with temozolomide. O^6^-methyl-guanine is both a cytotoxic and a mutagenic substrate. By mispairing with thymine, O^6^-methyl-guanine induces a G to A transition if the mispair is not corrected by the mismatch repair (MMR). This ubiquitous repair process corrects non-Watson Crick paired duplexes (63). MMR involves an excision and resynthesis mechanism for correcting the DNA mismatch (64). When the quantity of O^6^-methyl-guanine exceeds the cellular capacity to complete this process, single-stranded DNA and O^6^-methyl-guanine: thymine mispairs accumulate. The former lesion induces cytotoxicity, while tolerance of the latter leads to mutagenesis. Thus, cancer evolution in response to temozolomide represents a dynamic balance between the fate of these two lesions (65, 66). Each MGMT molecule transfers the methyl moiety from O^6^-methyl-guanine into itself before the methylated MGMT is targeted for degradation (65), and the repair of O^6^-methyl-guanine by MGMT is stochiometric (67). Thus, MGMT expression level is a key determinant in this dynamic interplay. The variability in MGMT expression associated with miR-181d degradation broadens the range of this interplay, which increases intra-tumoral heterogeneity.

The stability of most miRNAs (typically exceeding 12-24 hours) is primarily attributed to the binding of Argonaute (AGO), which protects the ends and the phosphate backbone of the miRNA from nucleolytic degradation (68, 69). While our understanding of miRNA degradation remains incomplete, the target-directed miRNA degradation (TDMD) mechanism proposes the binding of miRNA to a highly complementary sequence, termed “trigger RNAs,” induces a ternary complex that recruits E3 ubiquitin ligase, and an E2 ubiquitin-conjugating enzyme. This process results in the proteasomal processing of AGO, releasing the deprotected miRNA for nucleolytic degradation (70, 71). Whether and how the roles of ATR and PNPT1 in miR-181d degradation interface with TDMD remains an intriguing area of future research. It should be noted that the data presented in this study places ATR and PNPT1 only in a general pathway framework. For instance, the affinity-purified PNTP1 is limited in purity, and PNPT1 interacting proteins in the affinity pull-down may exert a more proximal effect on miR-181d. Moreover, our degradation assays were performed in the absence of AGO.

A framework that emerged from this study is that temozolomide exposure initiates a race between glioblastoma kill and acquired resistance for MGMT expression glioblastomas. The relative kinetics of these two processes ultimately determine the clinical outcome. The feed-forward loop between miR-181d and PNPT1 leads to rapid degradation of miR-181d, increasing 1) mean MGMT expression and 2) the variance of MGMT expression. The increased variation in MGMT expression is expected to facilitate the mutagenesis-related intra-tumoral heterogeneity, which enhances populational fitness to selective pressures related to future therapeutics. This framework offers one explanation for the poor prognosis associated with patients afflicted with MGMT-expressing glioblastomas (27, 72). Our study suggests that therapeutic delivery of miR-181d combined with TMZ should suppress the mean expression of MGMT and the variance of this expression, tilting the race towards tumor kill and clinical efficacy. This strategy offers promise where other therapies have failed for MGMT-expressing glioblastomas (27). It’s important to note that this approach is unlikely to apply to glioblastomas lacking MGMT expression, underscoring the need for genotype-specific therapies in future clinical translations.

## Materials and Methods

### Cell lines, cell culture, and plasmids

Human glioblastoma cell lines CMK3 (42) and BT-83 (41) were grown as neurosphere in NeuroCult media supplemented with heparin, human epidermal growth factor (EGF), human fibroblast growth factor (FGF) in ultra-low attachment flasks and kept at 37 °C incubator with 5% CO_2_.

Stable cell lines constitutively expressing miR-181d were established by transfecting pCMV-miR-181d miRNA expression vector (OriGene) into CMK3 and BT-83 cells selected with neomycin as described previously (26). Single clones were isolated and named CMK3 (miR181d), and BT-83 (miR181d), respectively. The corresponding negative control cell lines were generated by transfecting an empty control vector (OriGene) into CMK3 or BT-83 cells and selected with neomycin. PNPT1 knockout (KO) line was generated by transduction of PNPT1 gRNA containing pLenti-C-Myc-DDK-P2A-Puro lentivirus (OriGene) to CMK3 cells and selection with puromycin. The single KO clones were isolated and confirmed by DNA sequencing. A negative control cell line was generated by transfecting the corresponding empty expression vector.

### RNA isolation and quantitative RT-PCR (RT-qPCR)

Total RNA was isolated from cells using miRNeasy Kit (Qiagen) following the manufacturer’s protocol. cDNA was synthesized using miScript II RT Kit (Qiagen). miRNA and mRNA transcripts were quantified using SYBR Green (Bio-Rad) and target-specific primers (Qiagen) on the Bio-Rad Chromo 4 DNA Engine Thermal Cycler.

Newly diagnosed or and at the time of recurrence from the same patients’ glioblastoma surgical specimens were subjected to RNA isolation, reverse-transcribed, and subjected to qPCR of miR-181d, miR-939-5p, miR-3689d, miR-382-3p, miR-4513, miR-4519, miR-124-3p, miR-4323, miR-6657-3p, miR-6867-3p, miR-548aq-3p, miR-8071 and miR-5589-3p with miRCURY LNA primers (Qiagen) according to the manufacturer’s instructions.

### miRNA, siRNA, or plasmid transfection

CMK3 cells were transfected with human miR-181d (Qiagen, MIMAT0026608) and miR-21 mimic (MIMAT0000076), Bi-miR-181d (Qiagen, MIMAT0002821) or the non-targeting (NT) control (Qiagen, MIMAT0000010) using HiPerfect transfection reagent (Qiagen) following the manufacturer’s instructions.

siRNA transfections were carried out by HiPerfect transfection reagent (Qiagen). ON-TARGETplus siRNAs (Dharmacon) targeting human genes were used: RRP41 or ExoSC4 (L-013760-00-0010), PNPT1 (L-019454-01-0010), XRN1 (L-013754-01-0010), and non-targeting control (D-001810-01-10).

CMK3 PNPT1 KO cells were transfected with the pCMV6-AC-GFP vector (OriGene) encoding human WT-PNPT1 or point mutants (RPH1 domain: D135G; RPH2 domain: Q387R, R445E, R446E, R484A and catalytic sites: D538A and D544G) or empty vector by lipofectamine 2000 (Invitrogen).

### RNA sequencing (RNA seq)

BT-83 or CMK3 cells were treated with 500 µM TMZ or DMSO (1%) for 6 h. The cells were harvested, and RNA was isolated (miRNeasy Kit, Qiagen). The RNA quality was determined by Bioanalyzer (Agilent). The RNA samples were submitted to the University of Minnesota Genomic Center (UMGC) for library preparation and short-read sequencing.

### Clinical glioblastoma specimen collection and immunohistochemistry (IHC)

The research protocol (STUDY00012599) was approved by the Institutional Review Board of the University of Minnesota. All the patients gave informed consent before the study commenced. Only specimens that harbored wtIDH and umMGMT were collected for this study. Newly diagnosed or at the time of recurrence from the same patients’ glioblastoma surgical specimens were formalin-fixed and paraffin-embedded (FFPE). The FFPE sections were mounted on the slides and stained with MGMT antibody or Hematoxylin and Eosin (H&E). For MGMT detection, 4 μm sections were placed onto positively charged glass slides. The slides were deparaffinized and rehydrated. Antigen retrieval was performed. Endogenous peroxidase was blocked using 3% hydrogen peroxide, followed by Dako Protein Serum Block. Primary MGMT antibody (Abcam cat# ab39253, 1:100 dilution) was incubated for 30 min at RT. Detection was achieved using Rabbit EnVision™+ Kit (cat# K4003, Dako) and developed using diaminobenzidine (Dako) chromogen. Slides were counterstained with Mayer’s Hematoxylin. The IHC sections were evaluated and scored based on a previously published scoring system (73). In brief, for each MGMT stained section, ten random 400X fields were evaluated (5 random intra-tumoral and five random peri-tumoral fields). In each field, the density of MGMT positive cells was scored as No (no MGMT positive cells), Low (MGMT positivity in 1-33% of nucleated cells), Intermediate (MGMT positivity in 34-66% of nucleated cells), High (MGMT positivity in 67-100% of nucleated cells).

### Limiting dilution neurosphere forming assays

CMK3 cells were seeded in a 12-well plate at 10,000 cells per well. The cells were transfected with siPNPT1 or siNT. Twenty-four hours post-transfection, the cells were treated with or without 500 µM TMZ for 12 h. The cells were collected, single-cell suspensions were prepared, serially diluted, and inoculated into 96-well ultra-low attachment plates at specific densities. Cells were allowed to grow for 3 weeks. Each well was then examined for the absence or presence of a neurosphere (at least one aggregate of 10 or more cells was counted as one sphere). The frequency of sphere-forming cells was then calculated using Extreme Limiting Dilution Analysis (ELDA, http://bioinf.wehi.edu.au/software/elda/).

### Immunoprecipitation, affinity purification, and Western blot analysis

Dynabeads Protein G (30 µL) (Invitrogen) were washed two times in 10-bed volume of IP lysis buffer (20 mM Tris-HCl [pH 7.4], 150 mM NaCl, 2 mM EDTA, 1% NP-40). The beads were incubated with anti-rabbit PNPT1 antibody (Proteintech, 14487-1-AP or Abcam, ab157109) or anti-mouse turbo GFP antiserum (Origene, TA150039) or anti-mouse IgG (Santa Cruz, sc-2025) in 10-bed volume of IP lysis buffer containing 1 mM BSA (Thermo Fisher) for 45 min at room temperature. The bead-antibody complexes were washed once in a 10-bed volume of IP wash buffer (20 mM Tris-HCl [pH 7.4], 300 mM NaCl, 0.5% NP-40) and incubated with 300 µg of total cell lysate or C-terminal GFP tagged WT-PNPT1 or point mutant PNPT1 expressing cell lysate at 4 °C for 2 h with rotation. The enriched immune complexes were washed four times in IP wash buffer and collected by boiling with 1X SDS sample buffer.

Dynabeads M-280 Streptavidin (25 µL) (Invitrogen, 11206D) were equilibrated with a 10-bed volume of 1X PBS containing 0.5% BSA. The beads were incubated with 300 µg lysate of the cells transfected with the biotinylated miRNA at 4 °C overnight with rotation. The beads were washed four times in IP wash buffer and collected by boiling them with 1 X SDS sample buffer.

Cell lysates or affinity-purified proteins collected in SDS-sample buffer were resolved in SDS/PAGE and transferred onto the nitrocellulose membranes (Bio-Rad). Membranes were blocked using 5% BSA for 1 h at room temperature, followed by primary antibody incubation overnight at 4 °C. After washing with 1 X TBST buffer, membranes were incubated with species-specific secondary antibodies for 1 h at room temperature and developed in chemiluminescence reagent (GE biosciences) using Bio-Rad ChemiDoc MP Imaging System.

### Luciferase reporter assay

Luciferase reporters bearing the 3’UTR of PNPT1 were constructed by subcloning the 3’UTRs into pSiCheck-2 dual luciferase vector (Promega), including full-length 3’UTR, truncated sections of 3’UTR containing predicted miR-181d miRNA Response Elements (MREs), or truncated sections of 3’UTR containing mutated miR-181d MREs. CMK3 cells were co-transfected with miR-181d mimic and 3’UTR-luciferase reporter using Lipofectamine 2000 (Invitrogen) following the manufacturer’s protocol. Luciferase activity was measured 48 h post-transfection.

### TMZ resistance clone generation

BT-83 (0.5 x10^6^) cells were seeded in a 6-well plate. The cells were treated with 500 µM TMZ or 1% DMSO for 48 h. Post-treatment, the cells were washed and replenished with fresh culture medium without TMZ. After 4 weeks, the live cells started forming colonies. The colonies were collected, single-cell suspension was prepared, serially diluted and inoculated into 96-well plates at one cell/ well densities. Cells were allowed to grow for 4 weeks. Each well was then examined for the single cell clone, and the isolated clones were quantified for MGMT, miR-181d expression.

### Xenografts development and survival studies

Animal studies were performed following the Guide for the Care and Use of Laboratory Animals (Guide for the Care and Use of Laboratory Animals, 8th edition National Research Council (US) Committee for the Update of the Guide for the Care and Use of Laboratory Animals. Washington (DC): National Academies Press (US); 2011. ISBN-13: 978-0-309-15400-0ISBN-10: 0-309-15400-6). The animal study protocol was approved by the Institutional Animal Care and Use Committee (IACUC Protocol 2206-40091A).

For intracranial xenograft experiments, patient-derived BT-83 glioblastoma cells stably expressing miR-181d or miR-empty controls were dissociated into single-cell suspensions and stereotactically injected into the brains of nude mice at 6-weeks of age using murine stereotaxic system (Stoelting Co). The coordinates used were: 1.8 mm to the right of the bregma and 3 mm deep from the dura. Following tumor implantation, mice were maintained until the onset of overt neurological symptoms, including weight loss, lethargy, and hunched posture. After 7-days of tumor implant, the mice were randomly assigned to treatment groups (10 mice/group), and 50 mg/kg/day TMZ was administrated intraperitoneally for 5-days followed by 23-days treatment interruption. The mice were maintained for up to 90-days and the Kaplan-Meier survival curve was calculated. p values were determined using the log-rank test.

### Statistical analysis

Three or more independent experiments were performed for each assay and results were combined to define the mean ± Standard Deviation (SD). Statistical analyses were conducted using GraphPad Prism software 10. The statistical significance was evaluated using unpaired two-tailed Student’s t-test or one-way ANOVA. P value (*) of ≤ 0.05 was considered statistically significant.

## Acknowledgements

We thank Valya Ramakrishnan for her valuable technical support and for conducting key preliminary experiments that contributed to this study. We are grateful for the assistance provided by and discussions with Jie Li, Brian Hirshman, Ming Li, Shan Zhu, and Jianfeng Ning.

