## Supporting Information for "Modulating populational variance of methyl-guanine methyl transferase expression through miR-181d degradation: a novel mechanism of temozolomide resistance"

Gatikrushna Singh, PhD

**This supporting information file includes:**

Figures S1 to S4

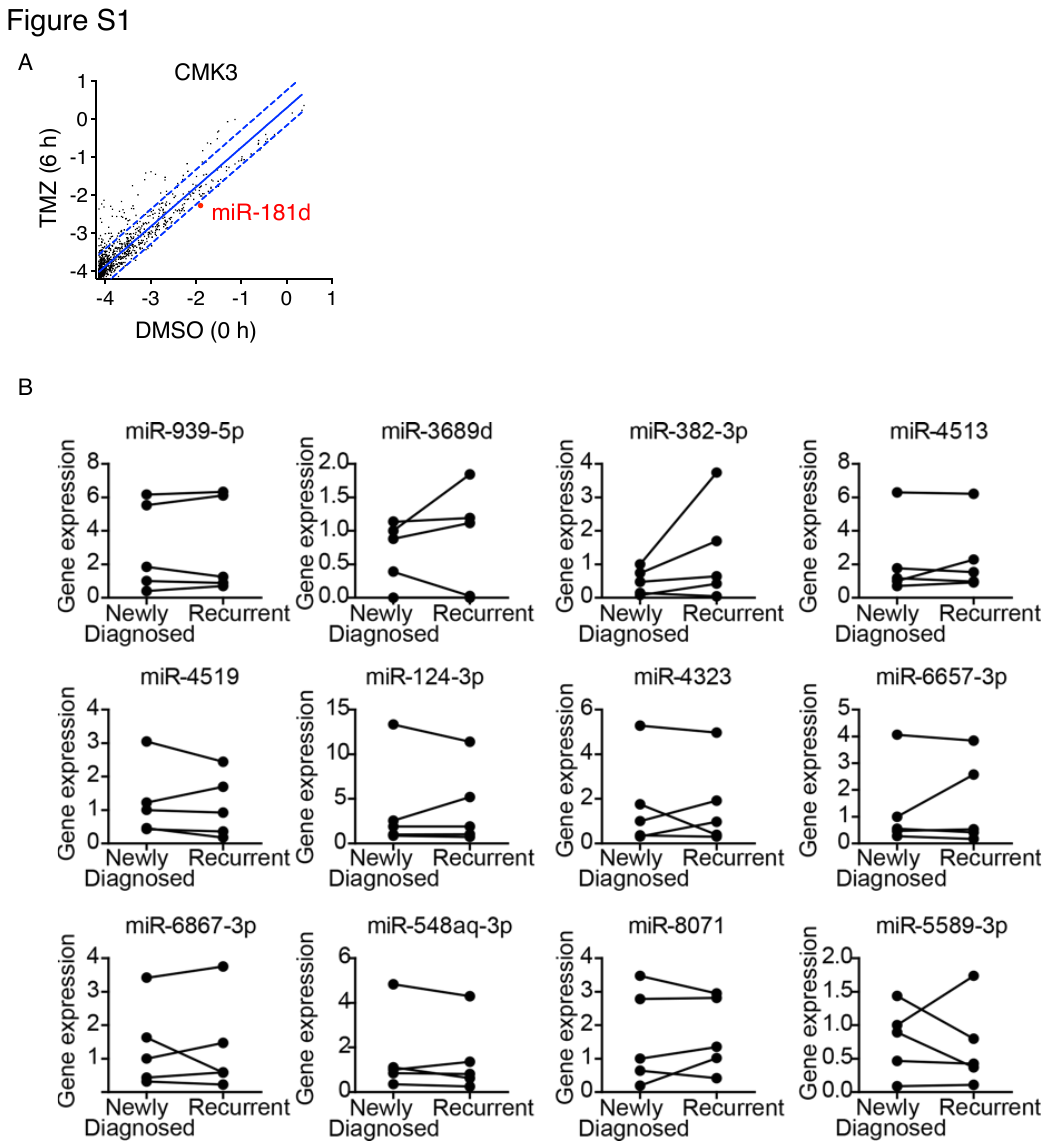

**Figure S1. TMZ treatment alters miRNA expression in glioblastoma cells.**

**(A)** Scatter plot representing the differential expression analysis of miRNAs from RNA-seq data of patient-derived CMK3 glioblastoma cells treated with 500 µM TMZ for 6 h. miRNA expression levels in TMZ-treated cells were compared to vehicle-treated cells (DMSO). miRNAs falling outside the diagonal lines indicate significant expression changes following TMZ treatment. miR-181d, which shows a notable reduction, is highlighted in red. **(B)** Quantification of miRNAs expression in 5 matched pairs of newly diagnosed (pre-TMZ treatment) and recurrent (post-TMZ treatment) formalin-fixed paraffin-embedded (FFPE) glioblastoma surgical specimen. miRNA expressions were measured using RT-qPCR.

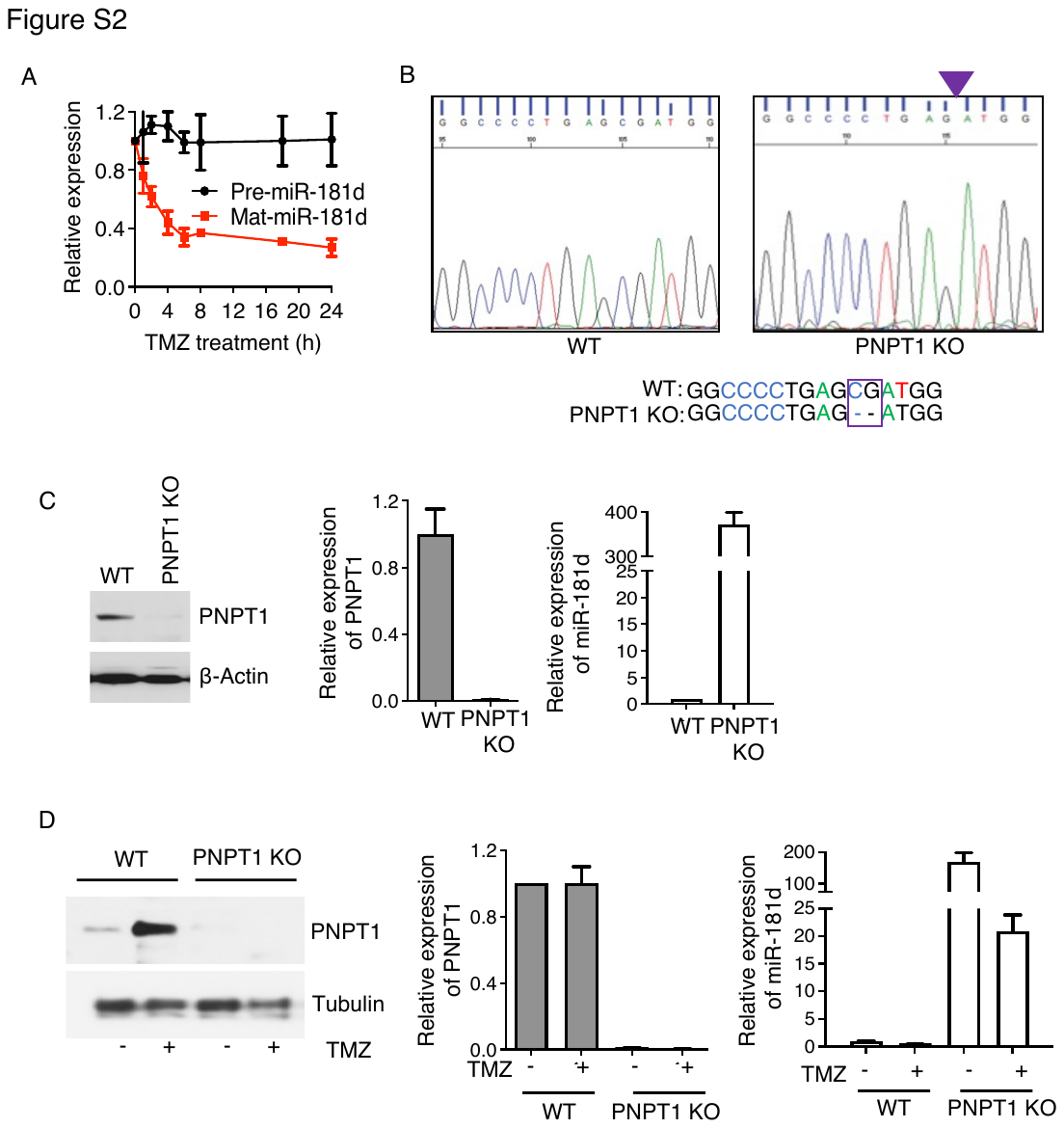

**Figure S2. TMZ enhances PNPT1 mediate-degradation of miR-181d.**

**(A)** BT-83 cells were treated with 500 µM TMZ up to 24 h. RNA was isolated, and precursor (pre) or mature (mat) miR-181d expression was analyzed by RT-qPCR. miRNA expression was normalized to vehicle-treated (DMSO) controls. **(B)** PNPT1 KO generation by CRISPR/Cas9 and disruption of PNPT1 confirmed by Sanger sequencing. **(C)** Western blotting and RT-qPCR analysis of PNPT1 and miR181d expression in PNPT1 KO cells. **(D)** PNPT1 KO cells knockout treated with TMZ. Western blotting and RT-qPCR analysis of PNPT1 and miR181d expression.

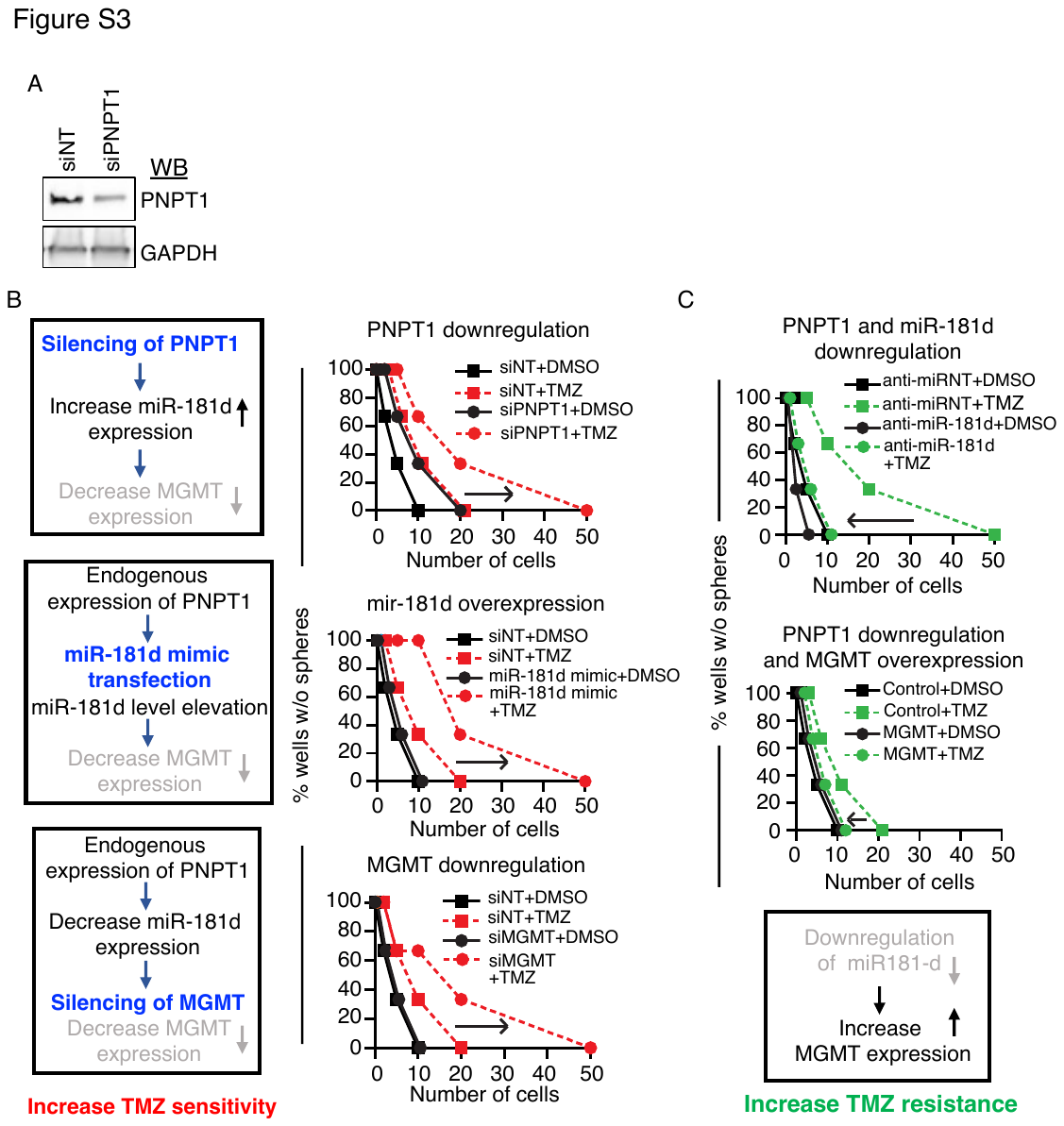

**Figure S3. PNPT1 regulates TMZ sensitivity by miR-181d and MGMT regulation.**

**(A)** CMK3 cells transfected with siPNPT1 or miR-181d or siMGMT and treated with TMZ (500 µM) or DMSO. The clonogenic potentials were determined by limiting dilution neurosphere forming assay. Summary of the assay represented in a flow chart as PNPT1/miR-181d/MGMT signaling to regulate TMZ sensitivity. **(B)** CMK3 cells were pre-treated with PNPT1 siRNA, before transfection with anti-miR-181d or MGMT-expressing plasmid (pCMV6-AC-GFP-MGMT). A limiting dilution neurosphere forming assay was performed, and a summary of the assay was represented in the flow chart.

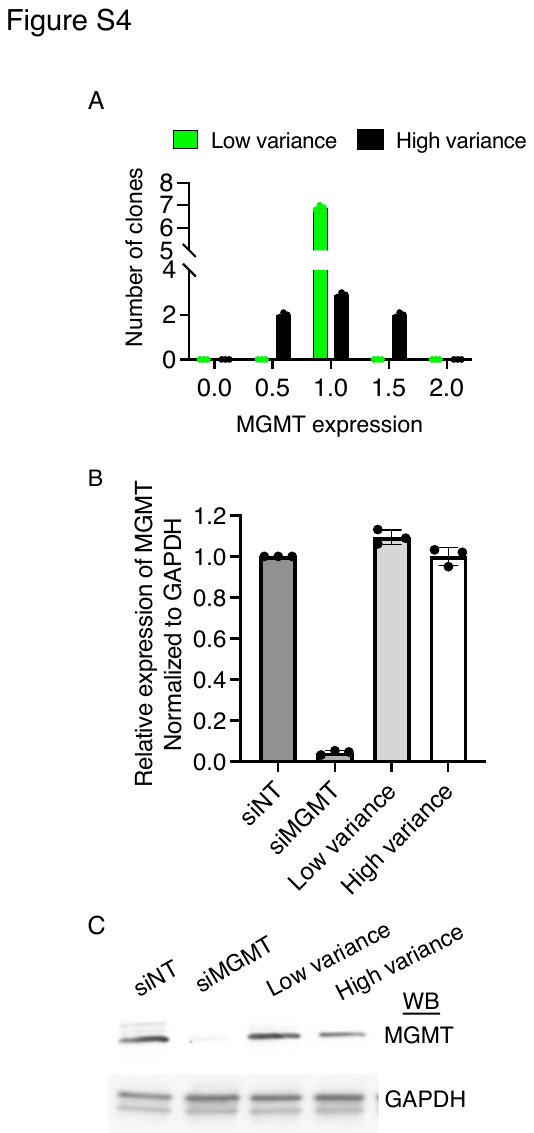

**Figure S4. Variability in MGMT expression.**

**(A)** Seven BT-83 clones expressing similar MGMT expression were pooled to generate either a low variance population (green) and high variance population (black) generated by combining three (comparable MGMT expression with low variance), two (higher MGMT expression) and two (lower MGMT expression) clones. MGMT expression was analyzed by RT-qPCR. **(B)** Quantitative RT-PCR analysis of MGMT mRNA expression in comparable low- and high-variance pools and MGMT siRNA transfected cells. **(C)** Western blotting of MGMT protein expression in the cells described in (B).
